# Predictions of Protein-Protein Interactions in *Schistosoma Mansoni*

**DOI:** 10.1101/233072

**Authors:** Javona White Bear, James H. McKerrow

**Affiliations:** Department of Bioengineering and Therapeutic Sciences, Department of Pharmaceutical Chemistry, and California Institute for Quantitative Biosciences, University of California, San Francisco, CA 94158; Graduate Group in Bioinformatics, University of California, San Francisco, CA 94158, USA; Department of Pathology and Sandler Center for Basic Research in Parasitic Diseases, University of California at San Francisco, San Francisco, California 94158, USA

## Abstract

**Background:** *Schistosoma mansoni* invasion of the human host involves a variety of cross-species protein-protein interactions. The pathogen expresses a diverse arsenal of proteins that facilitate the breach of physical and biochemical barriers present in skin, evasion of the immune system, and digestion of human hemoglobin, allowing schistosomes to reside in the host for years. However, only a small number of specific interactions between *S. mansoni* and human proteins have been identified. We present and apply a protocol that generates testable predictions of *S. mansoni*-human protein interactions.

**Methods:** In this study, we first predict *S. mansoni*-human protein interactions based on similarity to known protein complexes. Putative interactions were then scored and assessed using several contextual filters, including the use of annotation automatically derived from literature using a simple natural language processing methodology. Our method predicted 7 out of the 10 previously known cross-species interactions.

**Conclusions:** Several predictions that warrant experimental follow-up were presented and discussed, including interactions involving potential vaccine candidate antigens, protease inhibition, and immune evasion. The application framework provides an integrated methodology for investigation of host-pathogen interactions and an extensive source of orthogonal data for experimental analysis. We have made the predictions available online for community perusal.

**Author Summary:** The *S. mansoni* parasite is the etiological agent of the disease Schistomiasis. However, protein-protein interactions have been experimentally characterized that relate to pathogenesis and establishment of infection. As with many pathogens, the understanding of these interactions is a key component for the development of new vaccines. In this project, we have applied a computational whole-genome comparative approach to aid in the prediction of interactions between *S. mansoni* and human proteins and to identify important proteins involved in infection. The results of applying this method recapitulate several previously characterized interactions, as well as suggest additional ones as potential therapeutic targets.

## Introduction

### Etiological Agents and Effects of Schistomiasis

Schistosoma are dioecious parasitic trematodes (flukes) that cause the chronic disease Schistomiasis, affecting over 230 million people world-wide and causing more 200,000 deaths a year. They are digenetic organisms with six life cycle stages, four of which take place in the human host. [1] *Schistosoma mansoni*, one of the major etiological agents in Africa and South America of chronic Schistomiasis, releases eggs that become trapped in host tissues, triggering an unsuccessful immune response and causing a host granulomatous reaction, that in its most sever forms, causes fibrosis, scarring, and portal hypertension.

The host granulomatous reaction is a primary cause of mortality associated with Schistomiasis. [2, 3] Infection during childhood frequently results in growth retardation and anemia. The parasite persists in the host for up to 40 years with a high possibility of reinfection in endemic areas. [4] Standard methods for treating Schistosomiasis are not cost effective or affordable in most endemic populations, are ineffective as a prophylaxis against newly acquired infections (i.e. the cercarial and schisotsomula stages of the life cycle), and are becoming increasingly less effective against established infections due to parasite resistance. [5–7] Therefore, there is a need for improved and affordable treatments.[7,8]

### S. mansoni - Human Protein Interactions Involved in Pathogenesis and Infection Across Life Cycle Stages

*S. mansoni* infection involves parasite-human protein interactions over four of the six parasite life cycle stages [1]. Infection begins during the cercarial stage of the life cycle when the freshwater-dwelling larval cercariae contacts the human host. Invasion of the skin is achieved through degradation of the extracellular matrix by cercarial elastase; this same enzyme may help avoid the host immune response through cleavage of human C3 Complement [9] *S. mansoni* sheds its immunogenic tail to progress to the more mobile schistosomula life cycle stage, where it is carried by blood flow to the lungs and hepatic portal system.

Using proteomic analysis, cercarial elastase was implicated in the cleavage of an extensive list of human proteins, with follow-up experiments confirming its cleavage of at least seven dermal proteins [10]. After schistosomula entry, maturation to the adult life cycle stage occurs in the inferior mesenteric blood vessels where a number of proteins aid in immune evasion and digestion of human hemoglobin proteins. Among the proteins expressed in the adult cycle is the adult tegument surface protein Sm29, a potential Schistosomiasis vaccine candidate antigen. Sm29 interacts with unknown human immune proteins [11]. The final life cycle stage in humans is the egg phase; mated adults produce hundreds of eggs per day to facilitate transmission back to fresh water. The immune reaction to eggs leads to schistome pathogenesis. [12]

#### Large-Scale Computational Prediction of Protein-Protein Interactions

Ongoing efforts to address Schistomiasis include the development of new vaccines. Knowledge of the specific protein-protein interactions between the pathogen and human host can greatly facilitate this effort. However, a comprehensive literature review revealed only eleven confirmed interactions (Table 4), indicating the characterization effort is still in its infancy. These interactions were identified by experiments such as *in vitro* Edman degradation [10], fluorescence end point assay [13], crystallagrophy [14], and measurement of released radioactivity from a suspension [15]. Further types of low-throughput experiments could be based on hypothesis of specific predicted protein interactions [10].

While many methods have been developed to predict intraspecies protein-protein interactions, few have focused specifically on interspecies interactions, where knowledge of the biological context of pathogenesis can be used to refine predictions. Previously, our lab developed a protocol to predict interactions and applied these in the host-pathogen context [16,17]. Host-pathogen protein complexes were identified using comparative modeling based on a similarity to protein complexes with experimentally determined structures. The binding interfaces of the resulting models were assessed by a residue contact statistical potential, and filtered to retain the pairs known to be expressed in specific pathogen life cycle stages and human tissues (i.e. Biological Context Filter), thus increasing confidence in the predictions. The host-pathogen prediction protocol was benchmarked and applied to predict interactions between human and ten different pathogenic organisms [16,17].

#### Informing Computational Predictions

A crucial step for the construction of the Biological Context Filter is to annotate pathogen proteins by the life cycle stages in which they were expressed. This step is especially informative for *S. mansoni*, a digenetic organism, with life cycle stages and protein expression specific to both the molluscum and human hosts. While the human genome has been extensively annotated and made generally available, pathogen genome and expression annotation can be more elusive. Pathogens such as *Plasmodium falciparum* have been sequenced and extensively annotated [18]. In comparison, the *S. mansoni* genome, while much larger than that of *P. falciparum* (11,809 *S. mansoni* proteins vs 5,628 *P. falciparum* proteins), was only recently sequenced and assigned a full set of accession identifiers in GeneDB [19]. The sequencing effort is ongoing and there is limited annotation of corresponding proteins and structural information available. Thus, it is challenging even to cross-reference *S. mansoni* proteins described in various reports, particularly in those published prior to full genome sequencing [20]. Additionally, life cycle stage annotation is difficult to access or even absent in most databases.

The best source of annotation is directly from primary references in literature. Most primary reference accessions were often embedded in portable document format (pdf) files and other file formats that make extraction challenging. Furthermore, the context of an extracted accession and the life cycle stage to which it applies must be isolated from each reference and verifiable for accuracy and further study. We address these challenges by designing a simple natural language processing engine (NLP) that accomplishes data extraction, accuracy, verifiability, and correlation between disperse data sets required to construct the Biological Context Filter.

In Results below, we describe the benchmarking of our host-pathogen prediction protocol against experimentally characterized interactions, followed by a discussion of prospective prediction of novel host-pathogen interactions that may warrant experimental follow-up.

## Results & Discussion

### Protein Interaction Prediction

The protocol begins with the initial set of 3,052 *S. mansoni* and 8,784 human protein sequences (Fig. 1) for which high-quality models could be created using MODTIE [17]. The initial interaction predictions (Initial Predictions) were obtained by assessing host-pathogen interactions for which comparative models could be constructed. In previous work, the fraction of pathogen proteins that aligned to a protein template of previously observed solved complex structures averaged 21% [17]. Here, only 13.9% of *S. mansoni* proteins could be modeled using such a template. Human proteome interaction template coverage remained consistent with previous work at 34%. Overall, the protocol predicted 528,719 cross-species initial potential interactions. (Table 1).

**Figure 1.**
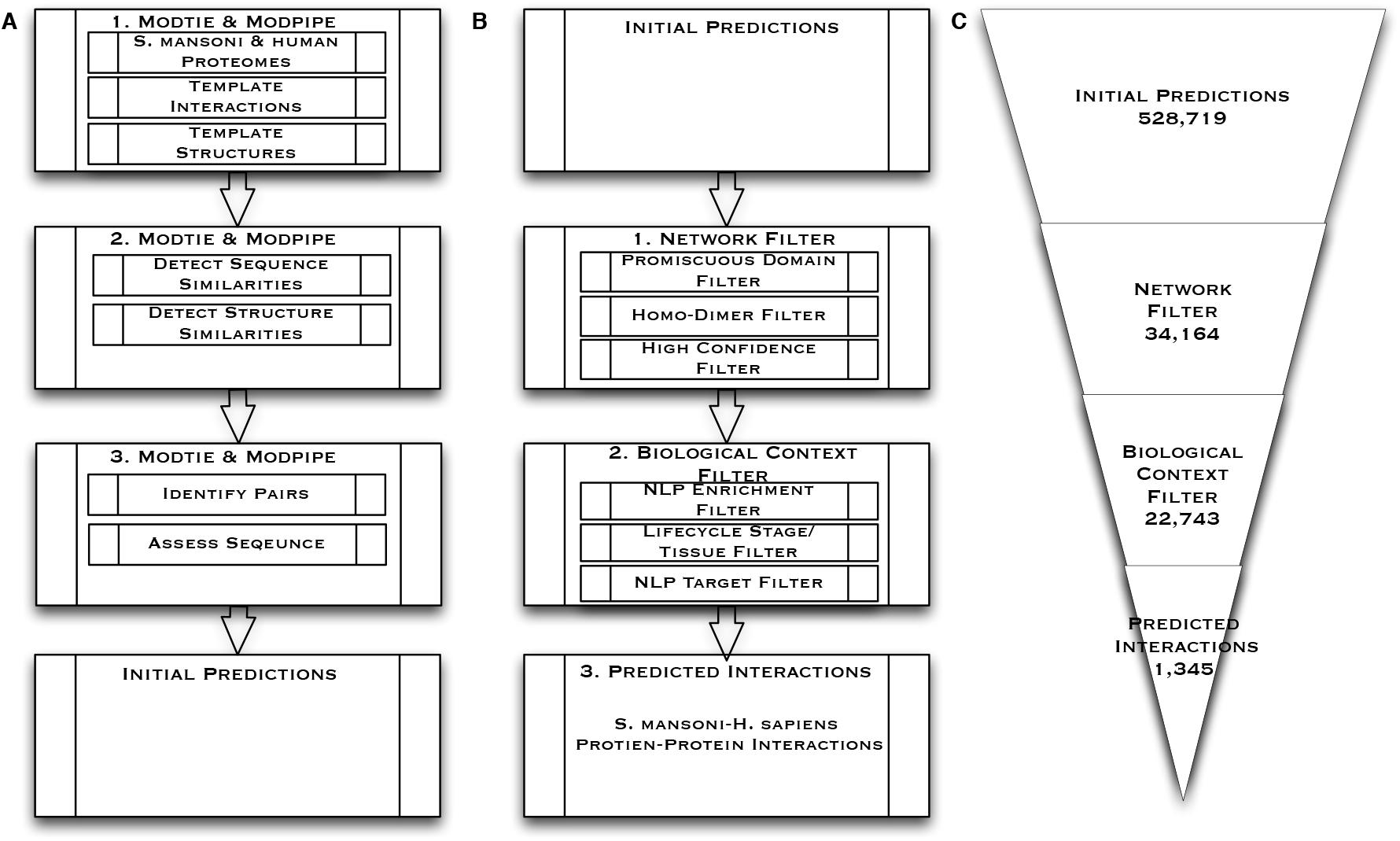
Prediction Framework: (A) Modtie & ModPipe protocol for detecting sequence and structure similarity: 1. The protocol begins with the set of human and *S. mansoni* proteins. 2. Sequence matching procedures are then used to identify similarities between the proteins and proteins with known structure or interactors. 3. A structure-based statistical potential assessment, or a sequence similarity score in the absence of structure, is then used to identify pairs with similarity to known complexes, assess the basis for a putative interaction, predict interacting partners and yield the Initial Predictions. (B) Initial Predictions: 1. Network Filter: Promiscuous Dimers, homo-dimer complexes, and high confidence interactions are then extracted from the initial set of predictions. 2. The Biological Context Filter is applied to the remaining set of predictions, weighing and ranking NLP enriched predictions, isolating life cycle stage and tissues interactions between *S. mansoni* and humans, and application of the Targeted Filter. 3. Predicted Interactions are an output of the Biological Context Filter. (B) Illustrates the reduction in interactions obtained after each step. The framework reduces the number of potential *S. mansoni*-human protein interactions by about three orders of magnitude (Table 1–3).

**Table 1.** Network Filter: Interactions removed during each application of indicated filter. Unfiltered interactions from the initial result set are shown in the first row. The number of interactions removed and remaining from each filtering step are shown in each category column including Promiscuous Pairs, Homo-dimer Complexes, and High Confidence interactions. The remaining 34,164 interactions are used as an input into the next step, the Biological Context Filter.

| Network Filter. |  |  |
| --- | --- | --- |
| Filter | Number of Interactions Removed | Interactions Remaining |
| Unfiltered | - | 528,719 |
| Promiscuous Domains | 242,677 | 286,042 |
| Homo-dimer Complexes | 143,065 | 142,977 |
| High Confidence | 108,813 | 34,164 |

Next, three network-level filters were applied to prune the initial predictions. The first Network Filter removed interactions based on promiscuous domains, defined as templates used for more than 1% of the total predictions. Promiscuous domains, while present in many interacting complexes lack specificity and are overrepresented in the predicted data set, making them less desirable as vaccine candidate antigens. For example, a domain in the crystal structure of HIV Capsid Protein (p24) bound to FAB13B5, Protein Data Bank (PDB 1E6J), is frequently used template in potential interactions.

However, immunoglobin is conserved in mammals and not specific to human interactions. Many variations will either not be applicable to *S. mansoni* and human interactions or conserved across species for binding similar epitopes. While most of the Fab (fragment antigen binding) region, which form the paratope, is highly variable in sequence, it is composed of less than 22 amino acids and relatively small. Many templates will score above the alignment threshold for this portion of the paratope and the short protein sequence acting as the epitope in the binding site based on shorter sequence lengths, high variability, and conservation across species resulting in a disproportionate number of potential interactions. Templates, similar to p24, were too generalized to draw conclusions with any degree of confidence and removed as promiscuous domains in the first Network Filter.

In the second Network Filter, predictions based on homodimer complexes were removed. This step removes instances where highly conserved interacting dimers form similar complexes in both *S. mansoni* and humans due to speciation events [21,22]. An example of such is the FGFR2 tyrosine kinase domain (PDB 1E6J). FGFR2 has high similarity to tyrosine kinases in *S. mansoni*, but were generally conserved in eukaryotes and thus comprise a bias in the homodimerization interaction of the catalytic subunits [23]. In the final Network Filter step, interactions with less than a 97% confidence interval were removed to further narrow the focus of potential interactions to a higher confidence level. This filtering results in the remaining 34,164 interactions.

Next, the Biological Context Filter isolates potential interactions according to various life cycle stages of *S. mansoni* and likelihood of in vivo occurrence in human tissues. In the first step of the Biological Context Filter, interactions passing the Network Filter were enriched with data from a simple natural language processing (NLP) literature search and databases using listed nomenclature and functional information (Methods). However, there is limited life cycle stage expression information using database annotation.

An NLP engine was designed to address this limitation and accomplished the following: (1) characterization of 12,720 *S. mansoni* genes automatically from primary reference; (2) recording of contextual, life cycle stage, and citation information into a customized database; and (3) programmatic correlation of this data with existing database annotations. This resulted in annotation of 96.6% of *S.mansoni* sequences, greatly exceeding existing annotation from any single database, which topped out at 62%. NLP annotation further extended this coverage with life cycle stage and characterization information not readily available in database annotation.

Next, the Life Cycle / Tissue Filter refines interactions for likelihood of in vivo interaction based on biological context derived from NLP of the component proteins in each interacting complex and their expression in the four life-cycle stages of *S. mansoni* different human tissues (Materials & Methods). The list of pathogen life cycle stage and human tissue pairs was generated (Table 2).

**Table 2.** Biological Context Filter: Interactions removed during each application of indicated filter. Unfiltered interactions from the Network Filter result set are shown in the first row. The number of interactions found from the total resulting data set are shown in each category column.

| Biological Context Filter Interactions. |  |  |
| --- | --- | --- |
| Filter | Interactions Removed | Interactions Remaining |
| Network | - | 34,164 |
| NLP & Enrichment | 10,929 | 23,235 |
| Life cycle/ Tissue | 492 | 22,743 |
| Targeted | 21,398 | 1,345 |

The third Biological Context Filter applies a targeted post-process analysis of potential interactions. In this step, NLP parameters were used to rank the prediction based on number of occurrences and the assigned weight of the literature where the observation occurred (Materials & Methods).

In previous work predicting host-pathogen protein interactions, filters resulted in a wide range of reductions for different pathogen genome due to varying levels of biological annotation available for each genomes. The majority of the biological annotations in Davis et. al. 2007 [17] were not relevant in a pathogenic context and therefore did not pass the filtering, while pathogen proteins had limited life cycle stage annotation resulting in multiple host-pathogen data sets with no interactions [17].

In the current framework, 22,743 (Table 2) interactions passed both biological and network-level filters, which was 51.5% more than the average of the ten pathogens in the previous work despite a below average model coverage. This increase is largely due to the NLP annotation, which produced a large number of pathogen proteins with a defined life cycle stage. Overall, the Biological Context Filter resulted in 1,345 annotated interactions in likely *in vivo S. mansoni* life cycle stage and human tissue interaction sites (Table 2).

#### Assessment I: Comparison of predicted and known S. mansoni-human protein interactions

To assess the predictions, we first compared the predicted set with the set of known *S. mansoni*-human protein interactions. There were 10 confirmed interactions between *S. mansoni* and human proteins. Among the 10, there is only one structure available in PDB (Table 4). The host-pathogen application framework recovered 7 of the 10 known interactions. The majority (7/10) of experimentally characterized *S. mansoni*-human protein interactions involve the serine peptidase cercarial elastase). Several experiments have characterized the cleavage by cercarial elastase of extracellular membrane and complement proteins [10,12,24–26].

Our method recapitulated several of these interactions. First, a retrospective prediction was made between the enzyme and human collagen based on the template structure of tick tryptase inhibitor in complex with bovine trypsin (PDB 2UUY) (Fig 2). Previous studies indicate that the enzyme has a role in suppressing host immune response (Table 3) [10,27]; its similarity to tryptase, which has been used as a indicator of mast cell activations and an important mechanism of host defense against pathogens [28],is consistent with this suggested role of cercarial elastase in pathogenesis.

**Figure 2.**
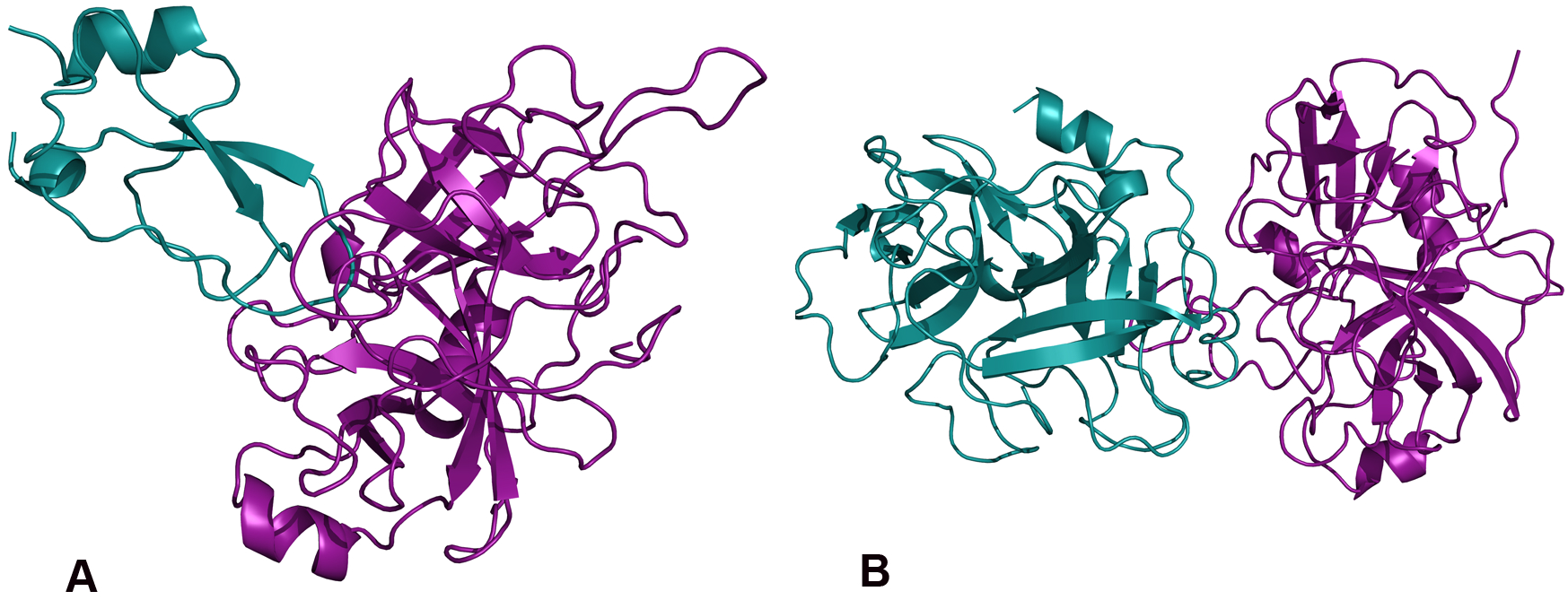
Retrospective Predictions: Examples of validated interactions. (A) Cercarial elastase (purple) and human collagen (blue) based on the template structure of tick tryptase inhibitor in complex with bovine trypsin (PDB 2UUY) (B) Cercarial elastase (purple) and human Complement C3 (precursor C3b) (blue) based on the template structure (PDB 1EQ9) of fire ant chymotrypsin complexed with PMSF, an inhibitor. Figures were generated by PyMOL (http://www.pymol.org).

**Table 3.** Protein interactions involved in *S. mansoni* life cycle stages that are directly involved in human pathogenesis from the resulting targeted predictions are shown with targeted life cycle stage, the number of predicted interactions for the corresponding life cycle stage and associated human tissues involved in the interaction. Human tissue expression data were obtained from the GNF Tissue Atlas [49] and GO [52] functional annotation unless noted otherwise.

| Biological Context Filter Interactions and *S. mansoni* Life Cycle Stage. |  |  |
| --- | --- | --- |
| Life cycle stage | Correlated Interactions | Life cycle stage-Tissue |
| Cercariae | 460 (460/1345) | 1 (skin) |
| Schistosomula | 442 (442/1345) | 15 |
| Adult | 329 (329/1345) | 13 |
| Egg | 114 (114/1345) | 11 |

In addition to cercarial elastases cleavage of extracellular proteins, several important protein-protein interactions involved in *S. mansoni* immune evasion have been characterized, including its cleavage of Complement C3 (Table 3). Our method retrospectively predicted this interaction based on the structure of fire ant chymotrypsin in complex with the PMSF inhibitor (PDB 1EQ9) (Fig 2). Fire ant chymotrypsin, which is similar to elastases in many species, degrades proteins for digestion and is a known target for blocking growth from the ant larval stage to adult in ant-infested areas [29].

#### Assessment II: Comparison of Vaccine Candidate Antigens and S. mansoni-human protein interactions

Next we assessed predictions against experimentally characterized vaccine candidate antigens where the mechanism, specificity, and interacting human proteins were still undetermined. Currently, there were 9 *S. mansoni* proteins considered as vaccine candidate antigens and 5 protein groups viewed as vaccine candidate antigens. We predicted interactions with 5 of the current vaccine candidate antigens and all of the potential vaccine candidate antigens (Table 5).

We now describe two specific examples of predicted interactions involving *S. mansoni* protein vaccine candidate antigens that may warrant experimental follow-up; these interactions are consistent with literature hypotheses. As noted, cercarial elastase is known to cleave several human proteins (Table 4), and it is considered a vaccine candidate antigen due to its abundance in *S. mansoni* cercarial secretions. Functionally, it has been indicated as the primary means of pathogen entry across the human dermal barriers, the first stage of pathogenesis [24].

**Table 4.** Confirmed protein-protein interaction between S. mansoni and human proteins. The application framework predicted 7 of the 10 known interactions. PDB column indicates the template PDB structure used to predict the interaction.

| Comparison of known and predicted *S. mansoni* protein interactions. |  |  |  |  |
| --- | --- | --- | --- | --- |
| S. Mansoni Protein | Human Protein | Predicted | Reference | PDB |
| Smp_001500 EIF4E | EIF4E-binding protein 1 | No | [14] | 3HXG |
| SmCe Cercarial elastase | Collagen (I, IV, VIII) | Yes | [15, 24, 25] | 2CHA |
| SmCe Cercarial elastase | IgE | No | [13] | - |
| SmCe Cercarial elastase | Complement C3 (C3b) | Yes | [10] | 1EQ9 |
| SmCe Cercarial elastase | Laminin | Yes | [15, 24, 25] | 2CHA |
| SmCe Cercarial elastase | Fibronectin | Yes | [15, 24, 25] | 2CHA |
| SmCe Cercarial elastase | Keratin | Yes | [53] | - |
| SmCe Cercarial elastase | Elastin | Yes | [54, 55] | 1FON |
| SmCB2 Cathepsin B | Collagen (I) (nidogen) | Yes | [10] | 1STF |
| SmCB2 Cathepsin B | Complement C3 | No | [10] | - |

We predicted novel interactions between cercarial elastase, its isoforms and other human proteins including calpains, cystatins, tetraspanins, immune and complement proteins (Table 5). An example is the interaction between cercarial elastase and the elastase specific inhibitor elafin. This prediction is based on the template crystal structure of elafin complexed with porcine pancreatic elastase (PDB 1FLE) (Fig 3). Elafin plays a wound-healing role in the dermal immune response in humans and is an antimicrobial against other pathogens such as *Pseudomonas aeruginosa* and *Staphylococcus aureus* [30]. Elafin has been demonstrated to bind with high affinity to both human leukocyte elastase and porcine pancreatic elastase. If a similar affinity can be characterized for its binding to cercarial elastase, elafin could act as a barrier to *S. mansonis* ability to establish infection. Experimentally validated binding with cercarial elastase could show that increased concentrations of elafin may be sufficient to prevent onset of infection and act as an effective barrier against infection [31].

**Figure 3.**
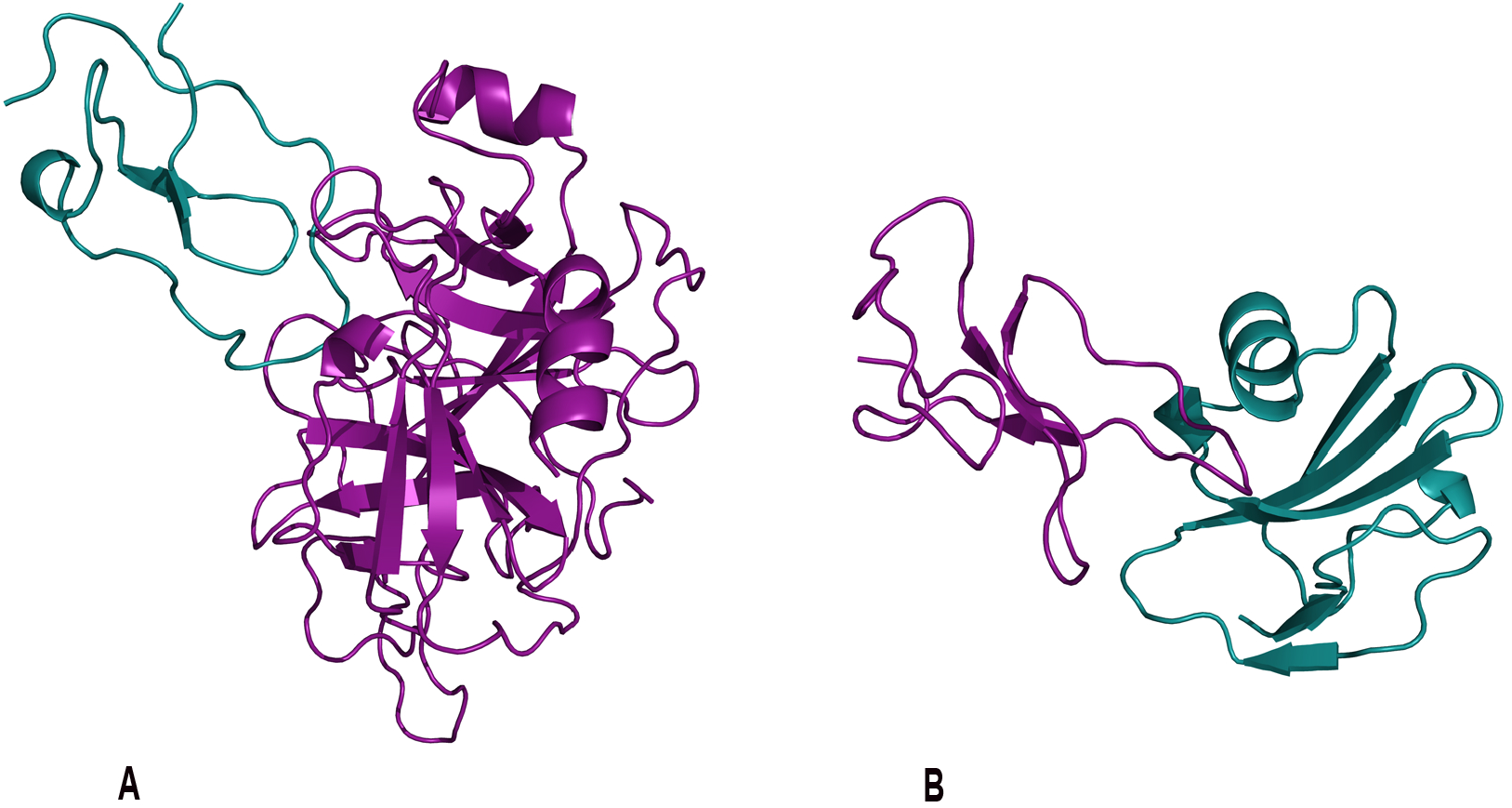
Prospective Predictions: Examples of predicted interactions. (A) Cercarial elastase (purple) was predicted to interact with the human elastase specific inhibitor elafin (blue). This prediction is based on the template crystal structure of elafin complexed with porcine pancreatic elastase (PDB 1FLE). (B) Sm29 (purple) was predicted to interact with human CD59 (blue) protein involved in the membrane attack complex corroborates hypothesis of *S. mansoni’s* ability to disable immune response. This prediction was based upon the template of ATF-urokinase and its receptor (PDB 2I9B). Sm29, an uncharacterized transmembrane protein, is an *S. mansoni* surface protein in both the schistosomula and adult life cycle stages that has been indicated in several immune response interactions making it an important vaccine candidate antigen. Figures were generated by PyMOL (http://www.pymol.org).

**Table 5.** Prospective protein-protein interactions between *S. mansoni* and human proteins. Prospective interactions are hypothesized, but have little or no experimental evidence, and are currently under investigation as candidate antigens or potential vaccine candidate antigens. The application framework predicted interactions between several proteins and suggested *S. mansoni* and human interactions with further interactions listed in Supplemental Tables (1–6).

| Potential *S. mansoni* Protein Interactions. |  |  |  |  |
| --- | --- | --- | --- | --- |
| *S. Mansoni* Protein | Human Protein | Predicted | Reference | PDB |
| Sm-TSP-1 Tetraspanin | IgG1/IgG3 Immune Response | No | [56] | - |
| Sm-TSP-2 Tetraspanin | IgG1/IgG3 Immune Response | No | [56] | - |
| Sm 29 Transmembrane | 67782326 (GI) TGF-beta receptor | Yes | [11, 56] | 2I9B, 1YWH |
| Sm 29 Transmembrane | 67782324 (GI) TGF, beta receptor II | Yes | [11, 56] | 2I9B, 1YWH |
| Sm 29 Transmembrane | 42716302 (GI) CD59 glycoprotein precursor, MAC | Yes | [11, 56] | 2I9B, 1YWH |
| Sm 29 Transmembrane | 9966907 (GI) SLURP-1 | Yes | [11, 56] | 2I9B, 1YWH |
| Sm 29 Transmembrane | 4505865 (GI) PLAU | Yes | [11, 56] | 2I9B, 1YWH |
| Sm 29 Transmembrane | 53829381 (GI) PLAU | Yes | [11, 56] | 2I9B, 1YWH |
| Sm 29 Transmembrane | 53829379 (GI) PLAU | Yes | [11, 56] | 2I9B, 1YWH |
| Sm 29 Transmembrane | 4504033 (GI) GPI anchored molecule-like | Yes | [11, 56] | 2I9B, 1YWH |
| Sm 23 Tetraspanin | IgG3, MAP-3 Immune Response | No | [11, 56–58] | - |
| Sm 14 FABP | IgG1, IgG3 Immune Response | No | [11, 26, 56, 58–62] | - |
| Sm 97 Paramyosin | IgG, IgE Immune Response | Yes | [58, 61, 63] | - |
| Sm 28 GST | IL-5, IgG2, MAP-4 Immune Response | No | [11, 58, 61] | - |
| SOD SOD [Cu-Zn], Cytosolic | 4507149 (GI) SOD1 [Cu-Zn] | Yes | [11, 58, 59, 64] | 2AF2, 1JK9 |
| SOD SOD [Cu-Zn], Cytosolic | 118582275 (GI) SOD3 Extracellular | Yes | [11, 58, 59, 64] | 2AF2 |
| SOD SOD [Cu-Zn], Cytosolic | 4826665 (GI) Copper chaperone for SOD | Yes | [11, 58, 59, 64] | 2AF2, 1JK9 |
| Sm-p80 Katanin p80 WD40 | C3 Complement Immune Response | No | [56, 65] | - |
| Cercarial Elastase | 4505787 (GI) Elafin *Supplemental Table 1 | Yes | [31] | 1FLE |
| Venom Allergen Proteins (VAL) | *Supplemental Table 2 | Yes | [26, 56, 66] | * |
| Calpain | *Supplemental Table 3 | Yes | [11, 12, 58, 65, 66] | * |
| Cystatin | *Supplemental Table 4 | Yes | [58, 59] | * |
| Tetraspanin | *Supplemental Table 5 | Yes | [11, 56, 66, 67] | * |
| Immune evasion | Immunoglobulin Proteins *Supplemental Table 6 | Yes | [5, 11, 13, 32, 59, 68] | * |

The Schistosoma protein Sm29 another vaccine candidate antigen indicated in pathogenic immune evasion, was involved in several predictions. The predicted interaction with the human CD59 protein, involved in the complement membrane attack complex (MAC), would aid *S. mansoni*s ability to disable immune response. This prediction was based upon the template of ATF-urokinase and its receptor (PDB 2I9B) (Fig 3), which is involved in multiple patho-physiological processes. Sm29, an uncharacterized transmembrane protein, is an *S. mansoni* surface protein in both the schistosomula and adult life cycle stages that has been indicated in several immune response interactions, making it an important vaccine candidate antigen.

CD59, protectin, regulates complement, inhibits the membrane attack complex (MAC), prevents lysis and is exploited as an established immune evasion tactic used by viruses [32,33]. Murine experiments indicate immunization with rSm29 reduces *S. mansoni* parasite burdens and offers protective immunity; however, the exact mechanism has not been characterized. Experimental validation of the predicted interaction between Sm29 and CD59 could provide further insight on how *S. mansoni* inhibits the MAC and additional strategies for preventing this inhibition. Additional interactions involving vaccine candidate antigens and key targets are referenced in the supplement.

#### Limitations

The *S. mansoni* genome was only recently sequenced [20], and there were fewer than 39 validated crystal structures of pathogen proteins available, with only one of these in complex with a human protein. Initial predictions rely on sequence and structure comparison to known interacting complexes, thus the lack of available protein structures in complex limits the coverage of the protocol. Additional experimental efforts will increase coverage and accuracy, by identifying more *S. mansoni* and human protein interactions, more protein interactions in complex, and further characterizing the biology for comparative analysis.

Furthermore, template coverage is primarily restricted to domain-mediated interactions, although peptide-mediated interactions are also known to contribute to protein interaction networks [34]. Peptide motifs that mediate protein interactions were identified through a combination of computational and experimental methods [35,36], and application of these motif-based methods will likely expand the coverage of host-pathogen protein interactions.

#### Prediction Errors

Several factors affect the accuracy of the method. These include errors in the comparative modeling process [37], the coarse-grained nature of the statistical potential used to assess the interface residue contacts [16], and consideration of only interactions between individual domains that could lead to predicted interactions that were unfavorable in the context of the full-length proteins. Additionally, both *S. mansoni* and humans are eukaryotic species, which means core cellular components, such as translation machinery, metabolic enzymes, and ubiquitin-signaling components are conserved and comprise many of the initially predicted interactions.

We address the similarities in conserved structures using the Biological Context Filter to remove complexes where there was a low possibility of *in vivo* occurrence, homodimer complexes that clearly involve conserved machinery, and high frequency template domains that could indicate both conserved sequences and structures as well as sequence-structure bias due to lack of interacting template coverage. For example, *S. mansoni* has been shown to secrete chemokine binding proteins as a decoy mechanism that modulates the host immune response. These proteins would be difficult to identify and characterized using known proteomic analysis and would likely be homologous to human proteins and would introduce noise into the detection and isolation of these types of interactions [38].

#### Future Work

Computational prediction and identification of protein-protein interactions is an important aspect in the development of new vaccines and vaccine candidate antigens. A variety of approaches such as genomic proximity, gene fission/ fusion, phylogenetic tree similarity, gene co-occurrence, co-localization, co-expression, and other features that only make sense or are currently feasible in the context of a single genome [39]. Comparative approaches offer a broad spectrum analysis of protein-protein interactions based on previously observed interactions.

The results of the approach and targeted analysis used on *S. mansoni*-human protein interactions here suggest that sequence and structure-based methods are an applicable approach [16,40]. This method has several extensions including those that identify peptide motifs [34], sequence signatures [41] that mediate interactions, and further extension of analysis and interpretation strategies such as analysis of the genetic polymorphisms at loci encoding for the proposed interacting proteins.

#### Potential impact

We developed a computational whole-genome method to predict potential host-pathogen protein interactions between *S. mansoni* and humans. Our results show seven validated predictions already experimentally characterized and numerous potential interactions involving proteins indicated as vaccine candidate antigens or potential vaccine candidate antigens. Despite limitations in *S. mansoni* structural coverage, our results demonstrate that broad-spectrum data enrichment and analysis is an effective method for protein-protein interaction prediction and highlight several potential immunization targets against *S. mansoni*. A lists of high-confidence predictions from targeted analysis are available online at http://salilab.org/hostpathogen. Additionally, in the tradition of open source efforts of the biomedical scientific community, the application framework is available for download. In closing, we expect our method to complement experimental methods in and provide insight into the basic biology of *S. mansoni*-human protein interactions.

## Materials and Methods

The initial predictions of *S. mansoni*-human were generated based on a protocol described in [17], briefly reviewed here. First, genome-wide *S. mansoni* and human protein structure models were calculated by MODPIPE [42], an automated software pipeline for large-scale protein structure modeling [43]. MODPIPE uses MODELLER [44] to perform the canonical comparative modeling steps of fold assignment, target-template alignment, model construction, and model assessment. High-scoring models were deposited in MODBASE [45], a publicly accessible database of comparative models. Next, resulting models were aligned to SCOP domain sequences, and if a model aligned to a SCOP sequence with more than 70% identity, it was assigned that SCOP domain identifier. These annotations were used as the basis for a search in PIBASE, a database of domain-domain interactions. In this search, those models assigned a SCOP domain that was part of a PIBASE interaction were structurally aligned to the conformation of that domain in the complex. In cases where a human model was aligned to one domain in a PIBASE interaction and an *S. mansoni* model was aligned to the other domain, a putative modeled complex resulted. This complex was then assessed with the MODTIDE potential, which outputs a Z-score approximating the statistical likelihood of the individual domain interface residues forming a complex across the two proteins. A detailed description of the full protocol is available in [16]. We refer to the resulting set of predictions as Initial Predictions.

### Filtering Interactions

Two sets of filters were applied to the resulting interactions. The first filter, referred to as the Network Filter was based on aspects of the modeling and scoring process. The second filter, referred to as the Biological Context Filter, was based on the stages of the life cycle and tissue pairs (Figure 1).

### Application of Network Filters

Predictions based on templates used for more than 1% of the total number of *S. mansoni* and human interactions were considered promiscuous and removed. 242,677 (45.9%) (Table 1) interactions met this criterion due to the overall similarity in eukaryotic organisms for network level machinery [46] and to the lack of known structure information for *S. mansoni* proteins. High confidence interactions were isolated based on previous work demonstrating an optimal statistical potential Z-score threshold of -1.7, which gave true-positive and false-positive rates of 97% and 3%, respectively [17]. The homodimer complex filter removed predicted interactions based on template complexes formed by protein domains from the same SCOP family excluding highly conserved eukaryotic pathways. These predictions primarily consisted of multimeric enzyme complexes formed by host and pathogen proteins, as well as core cellular components such as ribosome subunits, proteasome subunits, and core cellular components [17]. In total, 143,065 homo-dimer complexes were removed from the filter set based on this criteria (Table 1).

### Application of Biological Context Filters

Interactions that pass the Network filtering are then filtered for biological context using the following methods.

#### Natural Language Processing (NLP) & Enrichment Filters

##### S. mansoni Protein Annotation

In preparation for applying the Life Cycle / Tissue Filter, a Natural Language Processing (NLP) protocol was created to automatically identify from the literature which *S. mansoni* proteins were expressed in different pathogen life cycle stages. *S. mansoni* protein database identifiers and their amino acid sequences were extracted from the GeneDB [19], National Center for Biotechnology Information [NCBI], TIGR [47], and Uniprot/TrEMBL databases [48]. A literature search identified experiments indicating proteins expressed in different *S. mansoni* life cycle stages and categorized each literature reference into corresponding life cycle stages. All literature was then mined using NLP to derive accessions and context information. Accessions were derived from the text with regular expression searches corresponding to the specifications of the database (for example, a word in the text matching the regular expression form [A-Z][0–9]5 indicates a Uniprot Accession).

Thus, for each paper, a list of protein accessions was obtained. All protein accessions were then mapped by comparing sequences to Smp accession, Uniprot accession, and NCBI accession, in that order of priority. Thus, the final result of NLP processing was a list of all accessions of proteins expressed in life cycle stages of *S. mansoni*. *S. mansoni* protein sequence data from these initial interactions were enriched from biological annotation obtained from MODBASE [45], GeneDb [19], NCBI, Uniprot [48], and primary reference in literature. The annotations included protein names, links to referenced resources, and any available functional annotation.

##### Human Protein Annotation

Human proteins were annotated for tissue expression (GNF Tissue Atlas [49], known expression on cell surface, and known immune system involvement (ENSEMBL [50]. Functional annotation for each protein was obtained from Gene Ontology Annotation (GOA) [51]. Human protein sequences were correlated with predicted interacting sequences to determine involvement [16].

#### Life Cycle Stage/ Tissue Filter

Next, the Biological Context Filter was applied to *S. mansoni* and human protein interactions in the four life cycle stages associated with pathogenesis and infections in humans. *S. mansoni* proteins were filtered by life cycle stage, known expression and excretion, using NLP and database annotation. An interaction had to be present in the host tissue associated with the specific stage of pathogenesis and that *S. mansoni* life cycle stage to be included in the resulting interactions. The following life cycle stage and tissue pairs were applied to filter interactions: (1) cercariae proteins and human proteins expressed in skin, (2) schistosomula proteins and human proteins expressed in skin, lungs, bronchial, liver, endothelial cells, immune cells, red blood cells, blood, T-cells, early erythroid cells, NK cells, myeloid cells, and B-cells, (3) adult *S. mansoni* and human proteins expressed in liver, endothelial cells, immune cells, red blood cells, blood, T-cells, early erythroid cells, NK cells, myeloid cells, and B-cells, and (4) eggs and human proteins expressed in liver, endothelial cells, immune cells, red blood cells, blood, T-cells, early erythroid cells, NK cells, myeloid cells, and B-cells.

#### Targeted Filter

The final step in the Biological Context Filter uses a targeted post process analysis based on NLP and database annotations using two additional data mining steps. For each of the interacting protein complex pairs, three parameters were analyzed: pairwise expression in both known human tissue target and *S. mansoni* life cycle stage as indicated by the Life Cycle / Tissue Filter, expression or involvement in known human immunogenic responses, and *S. mansoni* protein expression or involvement with human proteins targeted by other parasites.

Parameters for additional data mining in the target analysis include the following criteria: investigator-selected proteins of interest and NLP derived key terms that were used to target annotation data in protein names and functional annotation (Uniprot [48], GeneDB [19], Gene Ontology [GO] [52]). Proteins selected as targets were assigned weights composed of two factors: (1) an average weight of number of citations across all references to the number of actual references used and (2) an investigator-assigned rank (1-3) based on significance and scope of primary reference/ experiments of NLP sources. The names of proteins and functional annotation were mined for the weighted key terms. All investigator-selected proteins of interest were presumed to pass filter criteria and the remaining interactions were ordered based on key term weights, rank, and Z score.

### Assessments

Predictions were benchmarked against confirmed *S. mansoni*-human interactions, which were compiled from the literature. Prospective interactions were assessed using vaccine candidate antigens and hypothesized vaccine candidate antigens where interactions have not been confirmed although several potential human protein binders have been experimentally identified (Table 4). Orthogonal biological information implemented in the filters provided significant enrichment of observed interactions (97% of predicted complexes were enriched). The number of protein pairs was reduced by about three orders of magnitude and assessment against previously characterized interactions (63% of known interactions predicted) suggests the method was applicable for genome-wide predictions of protein complexes.

